# Case study: Digital spatial profiling of metastatic clear cell carcinoma reveals intra-tumor heterogeneity in epithelial-mesenchymal gradient

**DOI:** 10.1101/2021.05.27.445912

**Authors:** Duncan Yi-Te Wang, Tuan Zea Tan, Ya-Ting Tai, Jieru Ye, Wei-Chou Lin, Lin-Hung Wei, Ruby Yun-Ju Huang

## Abstract

Intrinsic intra-tumor heterogeneity (ITH) has been linked to worse patient outcomes. The development of spatial profiling technology has enabled the deciphering of ITH with multiple analysis readouts. Advanced ovarian clear cell carcinoma (OCCC), known to harbor ITH, is chemoresistant, poor prognostic, and possesses distinct molecular and histological characteristics. However, detailed spatial information of the nature of ITH within OCCC remains unclear. Here, we utilized the NanoString Digital spatial profiling (DSP) GeoMx platform to perform multiplex protein expression analysis on tumor samples of primary and colonic metastatic sites from one advanced OCCC patient. The spatial resolution revealed the existence of an epithelial-mesenchymal (EM) gradient within the metastatic tumor but not the primary tumor, and similar EM gradient was not observed within the primary tumor. The EM gradient exhibited a distinct pattern from the periphery to the core of the metastatic tumor. Compared to tumor cells at the primary site, there was an intermediate zone in between the tumor periphery and the tumor core in the colonic metastasis with differential expression patterns of pan-cytokeratin (PanCK), fibronectin (FN), smooth muscle actin (SMA), neural cell adhesion molecule (NCAM), integrin alpha X (ITGAX), and Ki-67. Our study provides the first spatially resolved *in situ* evidence of intermediate or hybrid EM states within the tumor samples of similar morphology. This not only demonstrates the promising applications of spatial profiling in precision medicine but also provides an unprecedented view of the EM gradient during the progression of cancer such as OCCC.

## Introduction

Intra-tumor heterogeneity (ITH), manifested by genetically and molecularly diverse tumor cells admixing with various composition of surrounding immune cells and stromal cells in the tumormicroenvironment (TME), is crucial in determining the therapeutic responses and patient outcomes ^[1, 2]^. The revelation of ITH has been made clearer with the development of technological platforms with high dimensional multiplexing capacity and single-cell resolution. Methods such as single-cell sequencing lose the spatial context, making the interpretation of the neighborhood effects of ITH difficult.

Nanostring Digital spatial profiling (DSP) preserves spatial information and permits multi-omics analysis of cellular heterogeneity across micro-niche compartments of a tissue. The benefits of DSP are multiplex readouts of proteins and RNAs, user-defined regions of interest (ROIs), and the ability to profile these analytes in fresh frozen or FFPE tissue while preserving spatial information. In brief, the concept about how DSP works is like IHC or ISH with advanced multiplexing technique. After the sample is deparaffinized and rehydrated, the antigen or target retrieval step will be implemented, and then the sample will be incubated with the mixture of visualization markers (VMs) and probes attached with oligonucleotides by photo-cleavable linkers, which are used for the targets of interest. Once the sample is fully scanned by DSP, users can select regions of interest (ROIs) and define the criteria of areas of illumination (AOIs) using various profiling modalities including geometric shapes, fluorescent signal, rare cell profiling, contour and gridded segmentation approaches. The DSP machine will then project UV light onto these well-defined AOIs, and the released oligonucleotides will be collected by a microcapillary system. Once the oligonucleotides are collected, they will be dispensed into a microtiter plate and counted via the nCounter system or NGS readout ^[3, 4]^. By the combination of fluorescent VMs, unique oligonucleotide-attached probes, and the oligonucleotide-barcode system, DSP achieve the goal of retaining the spatial information of the sample and acquiring multiplex readouts simultaneously.

Amongst all subtypes of epithelial ovarian carcinoma (EOC), which account for about 90% of ovarian carcinoma, ovarian clear cell carcinoma (OCCC) at advanced stage has long been known for its relatively poorer prognosis, worse outcome, and lower sensitivity to standard platinum-based chemotherapy compared to other subtypes of EOC ^[5–7]^. This is linked to the molecular and genetic heterogeneity in OCCC tumors ^[8, 9]^; however, detailed information of ITH in OCCC has not been reported at high spatial resolution yet. Further knowledge of the mechanisms and molecular events contributing to its aggressiveness is the key to the development of more appropriate OCCC treatment. Here, we report the workflow of the first spatial profiling study, as far as we know, of the primary and metastatic tumor samples from an advanced stage OCCC patient. Our result revealed a spatially correlated gradient in the distribution of epithelial- and mesenchymal-like features, which is observed in the metastatic sites but not the primary site, suggestive of clonal diversity during tumor progression.

## Results

### Case history

A 47-year-old woman with a history of endometriosis presented with progressive abdominal fullness and vaginal spotting. Ultrasound showed an abdominal mass with ascites. Tumor marker of CA-125 was elevated (1,174 U/mL). Computed tomography (CT) scan demonstrated a lobulated tumor of 22.8 cm in diameter located at the lower abdomen and the pelvic cavity, multiple omental and mesentery nodules, a hyper-vascular hepatic tumor at S7, and ascites. There was also a ground-glass opacity at the right hemithorax with right pleural effusion. She underwent suboptimal debulking surgery including total hysterectomy, bilateral salpingo-oophorectomy, infra-colic omentectomy, anterior resection, peritoneal tumor excision, and cytoreduction. Cytology from pleural effusions, ascites, and peritoneal washing was positive for malignant cells. Final pathology confirmed the diagnosis of clear cell carcinoma in bilateral ovaries, bladder, peritoneum, omentum, colon, and the separated leiomyoma. She subsequently received 7 cycles of tri-weekly paclitaxel (175 mg/m2) and carboplatin (AUC 5) plus 6 cycles of bevacizumab (200 mg, 4.87 mg/kg) since 2nd cycle of chemotherapy.

### DSP Workflow

The archival FFPE sections of the sample from the ovarian primary site (labeled as Pri) and the one from the colonic metastatic site (labeled as Met) from the patient were included (Fig. 1a). All the ROIs were selected and segmented based on referencing the H&E staining morphology and fluorescence signals side-by-side. Visualization Markers (VMs) used to discriminate between OCCC tumor cells and immune cells include nuclear stain Syto13 and fluorophore-conjugated antibodies against PanCK, a tumor cell marker, and CD45, a pan immune cell marker. Meanwhile, the 18-plex Nanostring Immune Cell Profiling Core was utilized to reflect protein expression levels. ROIs were subdivided into PanCK-positive segments and CD45-positive segments as AOIs. For Pri, 19 of PanCK-positive segments and 15 of CD45-positive segments were collected and passed QC; for Met, 21 of PanCK-positive segments and 2 of CD45-positive segments were collected and passed QC.

**Fig. 1:**
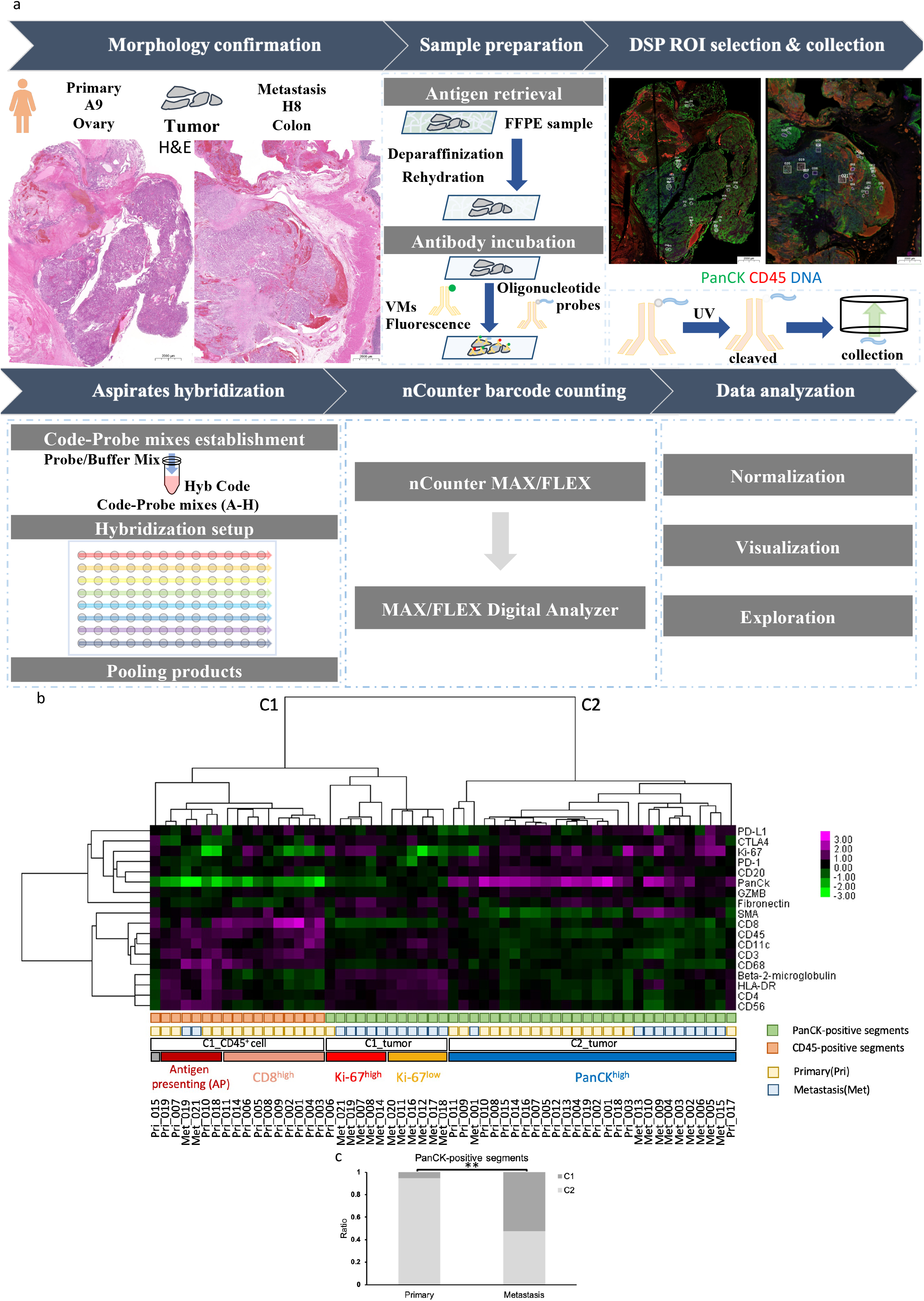
Digital spatial profiling (DSP) workflow. **a,** Workflow of DSP of the primary (Pri) and metastatic (Met) tumors. H&E-stained sections were used for morphology confirmation prior to sample preparation. Images of VM-stained fluorescence of DNA (blue), PanCK (green), and CD45 (red) were labeled with selected ROIs. Areas with wide range of bright red signals in Pri and dark orange signals in Met correspond to erythrocyte-rich regions indicating the occurrence of hemorrhage on H&E stains. H&E: hematoxylin and eosin. **b,** Unsupervised hierarchical clustering of the 18-plex protein expression data of PanCK-positive segments (grass green squares) and CD45-positive segments (orange squares) among all analyzed AOIs in the primary (light yellow) and metastasis (pale blue) tumors. Subclusters in colored squares from left to right: C1_CD45^+^ cell_Antigen presenting (dark red), C1_CD45^+^ cell_CD8^high^ (pale orange), C1_tumor_Ki-67^high^ (red), C1_tumor_Ki-67^low^ (yellow), and C2_tumor (blue). **c,** The ratio of the distribution of PanCK-positive segments within ROIs from the primary and metastatic tumors between clade 1 (C1, PanCK low and B2M high) and clade 2 (C2, PanCK high and B2M low), respectively. X-axis: tumor site. Y-axis: C1% vs. C2% ratio. Chi-square test, \*\**p* <0.01.

### Heterogeneity of PanCK-positive segments showed EM gradients in the metastatic tumor

The 57 segments were categorized into 2 clade and 5 subclusters (Fig. 1b). PanCK-positive segments, which represent tumor cells, could be divided into 2 clusters: C1_tumor and C2_tumor. These two clusters showed paradoxical expression patterns of PanCK and beta-2-microglobulin (B2M). For PanCK-positive segments in C1_tumor, low PanCK and high B2M were observed; for those in C2_tumor, high PanCK and low B2M were observed. All the PanCK-positive segments in C1_tumor were from Met except ROI 006 of Pri, while 64.3% of PanCK-positive segments in C2_tumor were from Pri. In contrast, 52.4% of PanCK-positive segments within Met belonged to C1_tumor, while the other 47.6% belonged to C2_tumor (*p*= 0.00371 by Chi-square test) (Fig. 1c).

Tumor cells in most of the PanCK-positive segments within Pri were, compared to those within Met, relatively more epithelial-like and homogenous. Almost all PanCK-positive segments within Pri belonged to C2_tumor (Fig. 1c). High homogeneity was observed in these PanCK-positive segments, and no certain distribution patterns of Pri tumor cells were observed (Fig. 2a).

**Fig. 2:**
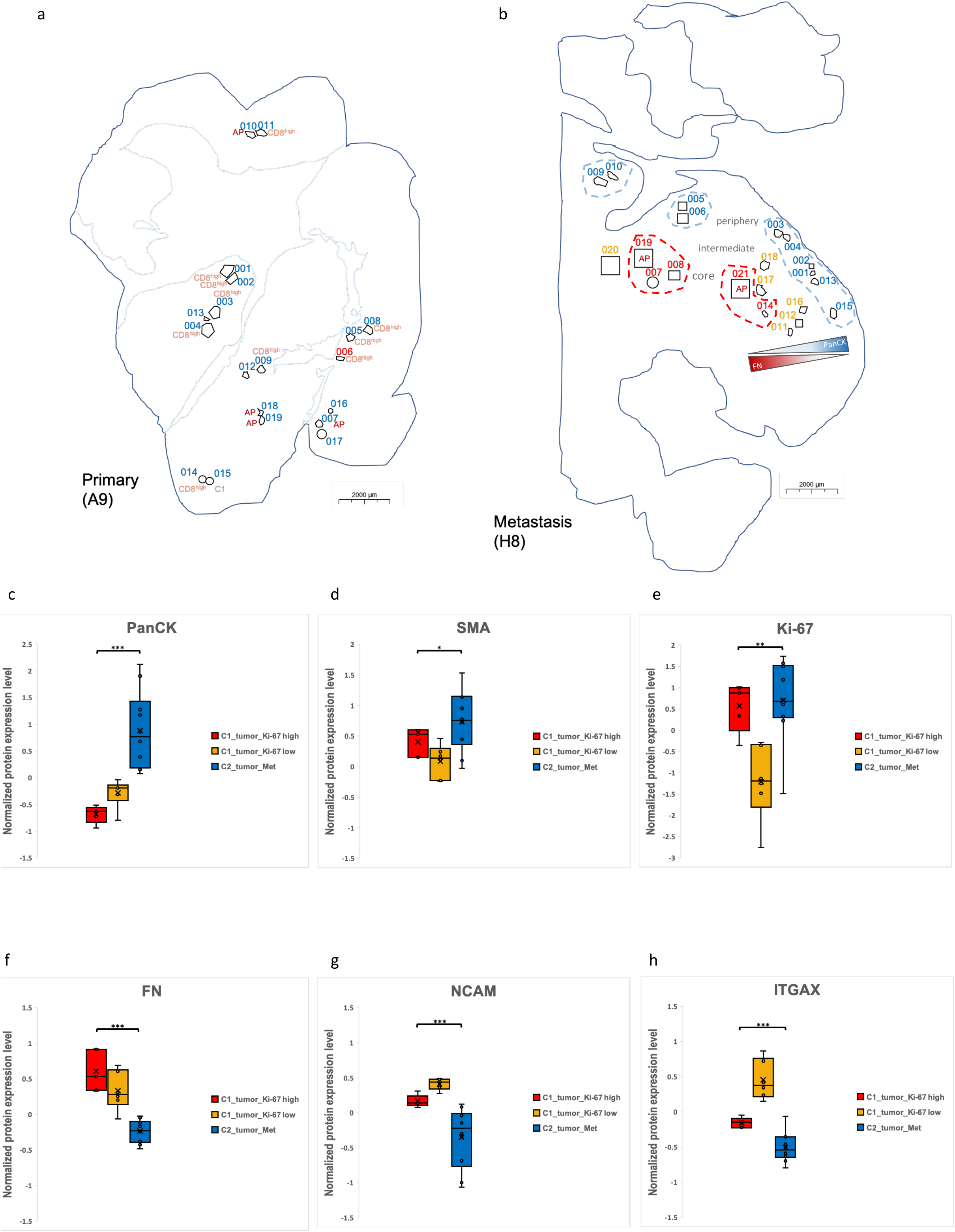
Spatial analysis of protein expression pattern showing an EM gradient. **a,** Outline of Pri morphology and selected ROIs. **b,** Outline of Met morphology and selected ROIs. Numeric labels of the ROI identity with corresponding colors indicate the subclusters of PanCK-positive segments. For CD45-positive segments within each ROI, additional text labels with corresponding colors were indicated next to the outline of ROIs. Scale bars indicate 2000 μm. **ch,** Box plots depicting the expression patterns of PanCK, SMA, Ki-67, FN, NCAM, and ITGAX between AOIs belonging to the C1_tumor_Ki-67^high^ (red), C1_tumor_Ki-67^low^ (yellow), and C2_tumor_Met (blue) subclusters in Met. Y-axis: numeric values of normalized protein expression level (extracted from the heatmap). One-way ANOVA, \**p* <0.05, \*\**p* <0.01, \*\*\**p* <0.001.

The data further suggested that tumor cells within Met were less epithelial and more heterogenous with lower expression of PanCK and high expression of SMA compared to Pri tumor cells. Met PanCK-positive segments belonged to two different clades: C1_tumor and C2_tumor_Met. Tumor cells in C1_tumor generally expressed lower level of PanCK and higher level of FN, B2M, HLA-DR, CD4, and CD56 compared to those in C2_tumor_Met, and tumor cells in C1_tumor could be further subdivided into C1_tumor_Ki-67^high^ and C1_tumor_Ki-67^low^ based on the expression level of Ki-67. Tumor cells in C1_tumor_Ki-67^high^ were relatively high in Ki-67, while those in C1_tumor_Ki-67^low^ were lower in Ki-67. Dissecting the spatial relationship between these segments within different clusters, we found that PanCK-positive segments of C2_tumor_Met were at the peripheral regions of Met, those of C1_tumor_Ki-67^high^ were at the tumor center, and those of C1_tumor_Ki-67^low^ were located in between (Fig. 2b). This spatial distribution pattern was corresponding to different hybrid EM (epithelial-mesenchymal) states. From C1_tumor_Ki-67^high^ (tumor center), C1_tumor_Ki-67^low^ (intermediate region) to C2_tumor_Met (tumor periphery), increasing levels of PanCK expression (Fig. 2c) accompany decreasing levels of FN (Fig. 2f) and CD56 (NCAM) expression (Fig. 2g), suggestive of an EM gradient. We, therefore, roughly classified C2_tumor_Met, C1_tumor_Ki-67^low^, and C1_tumor_Ki-67^high^ as early hybrid-E, early hybrid-M, and late hybrid-M according to the concept of EM Spectrum ^[10]^. Interestingly, tumor cells in C1_tumor_Ki-67^low^ expressed lower Ki-67 (Fig. 2e) and higher CD11c (ITGAX) (Fig. 2h) compared to those in C2_tumor_Met and C1_tumor_Ki-67^high^. Similar EM gradients were not observed in Pri.

We further performed geospatial analysis on PanCK-positive segments within Met to examine whether the changes in the expression of these markers correlated to the distance between these regions and the tumor core linearly. The results (Table 1) showed that the expression levels of PanCK (Fig. 3a) and SMA (Fig. 3b) were significantly positively correlated with the distance to the core; meanwhile, the expression levels of FN (Fig. 3c) and NCAM (Fig. 3d) were significantly negatively correlated with the distance to the core. Also, we found other 4 markers with linear correlation between their expression and the distance to core. These 4 markers included GZMB, B2M, CD3, and CD4 (Table 1). Among them, GZMB (Fig. 3e) had a positive correlation to the distance to core, while reverse situations were observed on B2M (Fig. 3f), CD3 (Fig. 3g), and CD4 (Fig. 3h). The geospatial analysis further confirmed the EM gradient within the Met tumor sample with significant anti-correlation between PanCK, SMA and FN, NCAM.

**Table 1:** The Pearson correlation coefficient of geospatial analysis on Met.

| **Protein** | **Pearson's r vs. distance to core (Mean)** | **p-value** |
| --- | --- | --- |
| PanCk | +0.6804 | 0.0004 |
| CD56 | -0.5801 | 0.0037 |
| Beta-2-microglobulin | -0.5575 | 0.0057 |
| Fibronectin | -0.5329 | 0.0088 |
| SMA | +0.4973 | 0.0158 |
| CD4 | -0.4863 | 0.0186 |
| CD3 | -0.4441 | 0.0338 |
| GZMB | +0.4379 | 0.0366 |
| CD20 | +0.3505 | 0.1011 |
| CD45 | -0.3361 | 0.1169 |
| CD11c | -0.3091 | 0.1513 |
| CD68 | -0.1882 | 0.3897 |
| PD-1 | +0.1523 | 0.4879 |
| PD-L1 | +0.1276 | 0.5616 |
| HLA-DR | +0.0490 | 0.8243 |
| CTLA4 | -0.0286 | 0.8970 |
| Ki-67 | +0.0117 | 0.9579 |
| CD8 | +0.0098 | 0.9646 |

**Fig. 3:**
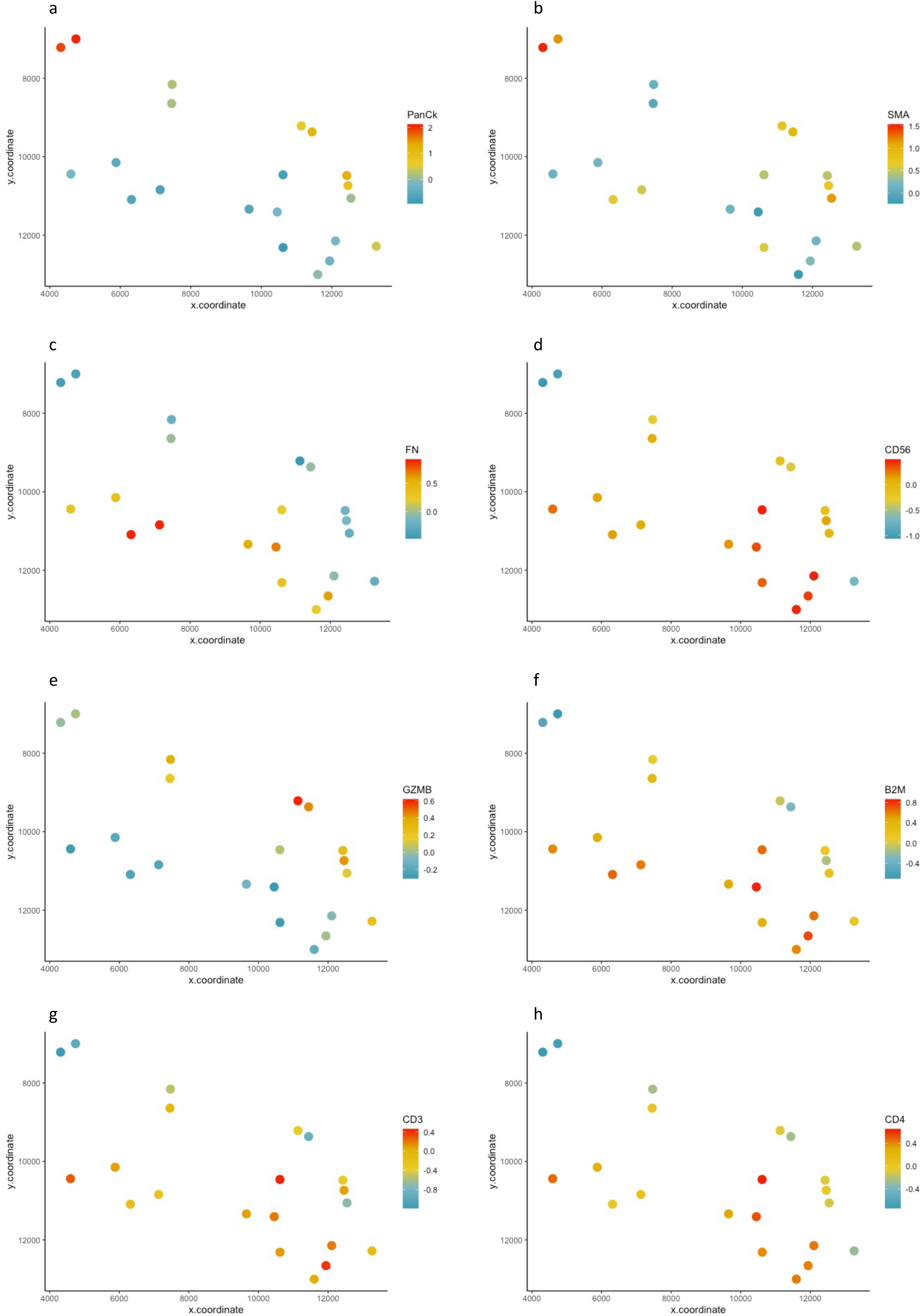
Geospatial analysis of the changes in the protein expression levels. **a-h,** Scatter plots depicting the changes in the expression levels of PanCK, SMA, FN (Fibronectin), NCAM (CD56), GZMB, B2M (Beta_2_microglobin), CD3, and CD4 between PanCK-positive AOIs of Met. The color of each point indicates the numeric values of normalized protein expression level (extracted from the heatmap). X-axis: x coordinate of the center of each ROI. Y-axis: y coordinate of the center of each ROI.

### Heterogeneity of CD45-positive segments

In general, CD45-positive segments in Pri and Met could be subdivided into two clusters (Fig. 1a), C1_CD45^+^ cell_Antigen presenting and C1_CD45^+^ cell_CD8^high^, by the expression of B2M, HLA-DR, CD4, and CD56. C1_CD45^+^ cell_CD8^high^, the cluster with relatively low expression of four immune-related markers, showed stronger CD8 signal than C1_CD45^+^ cell_Antigen presenting. There were no specific spatial distribution patterns observed within these two CD45-positive clusters, indicating that immune cell composition differs in different regions of the tumor.

## Discussion

OCCC has been shown to exhibit epithelial (E) and mesenchymal (M) subtypes based on the gene expression profiles ^[11]^. However, it is unclear whether there is intrinsic ITH of these EM gene expression subtypes within OCCC. In this study, we found that the EM heterogeneity shown as a gradient only exists in the metastatic but not the primary OCCC tumor. We speculated that these molecular changes and different EM states observed amongst these tumor cell subsets might be due to hypoxia in tumor.

It is widely accepted that the abnormal structure made by malignant cells limits the efficiency of oxygen diffusion and hinders oxygen delivery in solid tumor, and this thus makes the partial pressure of oxygen appear to be relatively low or even down to zero ^[12]^. Tumor hypoxia has been reported to correlate with several aspects of cancer such as poorer outcomes, angiogenesis, metastasis, genetic instability, tumor microenvironment, regulation of fibronectin and EMT ^[13–18]^. Indeed, nuclear expression of HIF-1α has been shown to express at significantly higher levels in OCCC than in other histological types (PMID: 17532031). In head and neck squamous cell carcinoma (HNSCC), a partial EMT (p-EMT) program has been shown to correlate strongly with the hypoxia program underlying the ITH of HNSCC at the leading edge of the tumor (PMID: 29198524). In our study, we speculated that comparatively low oxygen tension in tumor center caused environmental stress and initiated hypoxia-induced EMT in tumor cells near tumor center (C1_tumor_Ki-67^high^ & C1_tumor_Ki-67^low^), and this thus made them more mesenchymal-like than those at peripheral regions of tumor (C2_tumor_Met). This speculation could be further supported by the high FN and NCAM expression in C1_tumor_Ki-67^high^ and C1_tumor_Ki-67^low^. Studies have showed that hypoxia might lead to endogenous upregulation of FN, which could cause increased cell motility and activation of integrin-related signaling pathway ^[18–21]^, and relatively worse outcome of OCCC was already linked to higher expression of FN due to loss of p53 function previously ^[22]^. In the meantime, NCAM has been demonstrated to have association with migration and invasion of epithelial ovarian carcinoma ^[23]^, which strengthened the link between our observation of elevated NCAM expression in C1_tumor_Ki-67^high^ and C1_tumor_Ki-67^low^ and their mesenchymal-like properties. These studies indicate the importance of FN and NCAM in deciphering the nature of tumor cells in the metastasis tumor.

Intriguingly, upregulation of ITGAX and downregulation of Ki-67 in tumor cells of C1_tumor_Ki-67^low^ might reflect their aggressiveness. Although being long recognized as a dendritic cell marker, upregulated ITGAX has been reported to correlate with the invasiveness of breast cancer and prostate cancer ^[24, 25]^. Since ITGAX is a cell adhesion receptor molecule belonging to the integrin family, we speculated that its upregulation might involve in the activation of FN-integrin signaling pathway in cell migration. Meanwhile, the relatively low Ki-67 might reveal that tumor cells in C1_tumor_Ki-67^low^ were in quiescence. Ki-67 has been recognized as a cell proliferation marker, and cells with low Ki-67 expression are regarded as slow-growing cells or cells in G0 ^[26]^. Tumor cells entering mesenchymal-like states with slow proliferation rates might be one possible mechanism to escape from apoptotic and anoikis ^[27–29]^. Also, the low proliferation rate might associate with the trade-off between migration and proliferation ^[30]^. Previous research has demonstrated that ovarian carcinoma cells with intermediate mesenchymal state are more anoikis-resistant and aggressive compared to those with other EMT phenotype ^[31]^. Collectively, tumor cells in C1_tumor_Ki-67^low^ in an early-hybrid M state were equipped with some beneficial features against environmental stress.

EMT has been suggested to contribute to carcinoma progression ^[32]^. However, from the clinicopathological perspective, the EMT changes are often not visually discernable by routine histopathological examinations. Reasons being that EMT usually only occurs in a selected tumor cell population either at the invasive fronts or close to the hypoxic region ^[16]^. In addition, the detection methods for EMT in the clinicopathological settings have not been optimally established. Various EMT scoring scheme have been proposed ^[10]^, but none of these have yet to reach clinical use. The nature of tumoral heterogeneity is significantly reflected from our study which illustrates the spatial distribution of an EM gradient observed in the metastasis but not the primary tumors. This highlights the inter- and intra-tumoral heterogeneity which could not be resolved by traditional methods. The availability of technologies such as DSP has enabled the detection of the localized EMT changes that exists in morphologically similar tumor regions. This opens up a new avenue to explore the EMT spectrum in patient samples.

Though DSP is a powerful tool to investigate further into molecular expressions of cells in ROI, it was reported that the detection and measurement were influenced by the abundance of markers and the order of detection, which might result from sequential oligonucleotides collection step and intrinsic 10 μm resolution restriction ^[33, 34]^. From our experience, we observed that erythrocytes (diameter < 10 μm) in some ROIs would be mis-labelled as target cell in segmentation step, such as CD45-positive cells, and this could cause bias in data interpretation because of the interference of collected oligos from those erythrocytes. This can be prevented via elevating the threshold of DNA channel signal when doing segmentation or not selecting erythrocyte-rich areas as ROIs, but the interference would be sometimes unavoidable if the sample is, for example, from tumor section with hemorrhage, which would cause erythrocytes to overwhelm tumor cells in interested regions. Another limitation is that it is impracticable for DSP to have the entire tissue section analyzed currently. Compared to CODEX, for instance, DSP depends on the detection of several selected small ROIs (the maximum size of a ROI in diameter is about 760 μm) to acquire data utilized to conduct further exploration. Conceivably, it will be quite time and cost consuming to have the whole sample examined. Thus, beforehand investigation, appropriate study design with good research questions is indispensable to utilize the power within this high-throughput multiplex platform to the fullest.

## Methods

### Sample preparation

The study was approved by the Institution Review Board (IRB) of National Taiwan University Hospital (NTUH) (No. 202005080RIND). The patient provided written informed consent for the collection of tumor samples of this study. Formalin-fixed and paraffin-embedded (FFPE) tissue sections (5 μm) were baked at 60°C for 1 hour. Deparaffinization was performed using CitriSolv and sections were rehydrated in 100% and 95% ethanol sequentially and followed by wash in ddH_2_O. Antigen retrieval was performed by boiling at 121°C in pH 6 Citrate buffer solution for 15 minutes in a pressure cooker. Tissue sections were blocked in blocking buffer for 1 hour at room temperature before incubated with nanoString GeoMx DSP Immune Cell Profiling (a panel of antibodies with UV photocleavable oligonucleotide barcodes), PanCK-Cy3 (1:40) and CD45-Texas Red (1:40) antibodies overnight at 4°C in dark. Sections were stained in SYTO13 (1:10) for 15 minutes on the following day after post-fixed in 4% paraformaldehyde (PFA).

### ROI selection, collection, and analysis

Stained sections were loaded onto GeoMx DSP followed by the selection of region of interests (ROIs). All the OCCC samples in the study were annotated by a pathologist (W.C. Lin) on the H&E slides. ROIs selection was based on annotation of their respective H&E slides. The ROI size ranged from 200 μm to 700 μm in diameter, and 18 to 19 ROIs were selected per sample. ROIs can be compartmentalized into tumor or immune-related cell segments. Once ROIs and segments were confirmed, UV-light were projected onto each defined segment. UV photocleavable oligonucleotide barcodes were released, collected by microcapillaries, and dispensed into different wells of a microtiter plate, respectively. The collected oligonucleotide barcodes were then pooled and hybridized at 67°C for at least 16 hours before loading onto nCounter Flex system.

### Data normalization and visualization

The readouts from nCounter (version 4.0.0.3) were transferred to GeoMx, QC and normalization using the built-in data analysis software (version 2.1.0.33). The raw data were first normalized to ERCC (External RNA Control Consortium) spike-in controls. Subsequently, the spike-in normalized data were scaled by geometric mean of nuclei counts, and by geometric mean of Rb IgG. The normalized data were log-transformed, and genes, arrays were mean-centered by Cluster 3.0 ^[35]^. Clustering was also performed using Cluster 3.0 using similarity metric of Pearson correlation, and tree-construction method of centroid linkage. The heatmap of the clustered data was generated with Java Treeview (version 1.1.6r4) ^[36]^. The normalized data were extracted and made into box plots and scatter plots. The data of x coordinate and y coordinate utilized to build the scatter plots were extracted from DSP. The scatter plots were created in R using ggplot2 package. Statistical significance of association was assessed using *Chi-square* test whereas mean difference was assessed using Anova test.

### Distance to core analysis

Euclidean distance to the Met core was computed using Matlab® R2016b version 9.1.0.960167 (MathWorks; Natick, MA, USA) pdist2 function using the x- and y-coordinates of the ROIs. We estimated the core of Met from the x- and y-coordinates average from the 4 ROIs of the core.

## Data Availability

The data that support the findings of this study are available from the corresponding author on reasonable request.

## Disclosure Statement

The authors have no conflicts of interest to declare.

## Funding Sources

This work was supported by the Yushan Scholar Program by the Ministry of Education, Taiwan (NTU-110V0402) and NTU Core Consortiums (NTUCC-109L893001) to R.Y.-J.H; NTUCC-109L893003 to Lin-Hung Wei.

## Author Contributions

Conceptualization, R.Y.-J.H. and Lin-Hung Wei; writing, Duncan Yi-Te Wang, Ya-Ting Tai and R.Y.-J.H.; DSP data acquisition, Jieru Ye and Duncan Yi-Te Wang; data analysis and interpretation, Duncan Yi-Te Wang and Tuan Zea Tan; pathology review, Wei-Chou Lin; clinical review, Lin-Hung Wei and Ya-Ting Tai. All authors have read and agreed to the published version of the manuscript.

## Notes

### Competing Interest Statement

The authors have declared no competing interest.

